# Dissecting autonomous enzymatic variability in single cells

**DOI:** 10.1101/2024.10.03.616530

**Authors:** Christian Gnann, Alina Sigaeva, Trang Le, Anthony Cesnik, Sanem Sariyar, Diana Mahdessian, Rutger Schutten, Preethi Raghavan, Manuel D. Leonetti, Cecilia Lindskog, Mathias Uhlén, Ulrika Axelsson, Emma Lundberg

## Abstract

Metabolic enzymes perform life-sustaining functions in various cellular compartments. Anecdotally, metabolic activity is observed to vary between genetically identical cells, which impacts drug resistance, differentiation, and immune cell activation. However, no large-scale resource systematically reporting metabolic cellular heterogeneity exists. Here, we leverage imaging-based single-cell spatial proteomics to reveal the extent of non-genetic variability of the human enzymatic proteome, as a proxy for metabolic states. Nearly two fifths of enzymes exhibit cell-to-cell variable expression, and half localize to multiple cellular compartments. Metabolic heterogeneity arises largely autonomously of cell cycling, and individual cells reestablish these myriad metabolic phenotypes over several cell divisions. Multiplexed imaging revealed that metabolic states are continuous and that the correlation between metabolic pathways is metabolic state dependent. These results establish cell-to-cell enzymatic heterogeneity as an organizing principle of cell biology that may rewire our understanding of drug resistance, treatment design, and other aspects of medicine.

---

Cellular metabolism is an essential process that orchestrates the production of energy and building blocks for the cell, while also disposing of cellular waste. This is coordinated by a plethora of metabolic enzymes (including transporters), cofactors, and metabolites that form a complex dynamic network of chemical reactions. The reactions are tunable in response to intracellular or extracellular signaling, nutrient availability, cell cycle progression or circadian rhythm. Recent advances in single-cell sequencing technologies have revealed cell-to-cell heterogeneity in genetically identical cell populations.^1–3^ This heterogeneity results in phenotypic differences, including variable expression of metabolic enzymes that has been linked to cancer drug resistance^4,5^, metastasis^6^, differentiation^7,8^, and immune cell activation^9,10^.

Despite these insights, studies of cellular metabolism are mostly based on bulk proteomic or metabolomic measurements^1^, thereby limiting the ability to study metabolism with single-cell or subcellular resolution. Single-cell transcriptomics suffers from systematic noise and dropouts that make it difficult to study cellular heterogeneity.^11^ Additionally, although gene expression roughly predicts the expression of the corresponding protein, it fails to capture cellular variability that is established post-transcriptionally.^12^ Spatially resolved measurements of metabolites by matrix-assisted laser desorption ionization mass spectrometry imaging (MALDI-MSI) have revealed heterogeneity in the lipid composition in single cells.^12,13^ However, these studies observe a relatively small number of molecules and fail to capture the full complexity of metabolism on the single-cell level including its subcellular organization. Mass spectrometry (MS)-based subcellular proteomics can classically assess protein localization but does not provide single-cell resolution, rendering studies of single-cell heterogeneity impossible.^14^ Imaging-based subcellular proteomics using antibodies^15^ or fluorescently tagged proteins^16–18^ overcomes those limitations by simultaneously characterizing protein levels and distribution in individual cells with high resolution. Such subcellular measurements can reveal spatial or temporal protein expression heterogeneity,^14,19^ thus providing insights into the spatiotemporal partitioning of metabolic processes.

Here, we present the first global map of the metabolic proteome with single-cell and subcellular resolution. We show that more than half of all enzymes are multilocalizing (*i.e.,* localize to multiple cellular compartments). By integrating protein interactomic data with our imaging dataset, we dissect compartment specific interactions and identify novel multifunctional enzymes. Lastly, we show that metabolism exhibits a higher amount of heterogeneity than most other biological processes, with a comparable amplitude of variability to that of the cell cycle. Autonomous cell processes have been found in organelle biogenesis and cytoskeletal organization, with oscillations that are independent but influenced by the cell cycle^20^. We observe autonomy of metabolic heterogeneity from the cell cycle in this work and evidence that it is regulated mainly post-transcriptionally, resulting in a range of distinct metabolic states within one cell population. We believe that understanding these metabolic states is important to understanding human health and disease at a molecular level in fundamental, preclinical and clinical contexts. All data are publicly available as part of the Human Protein Atlas (HPA) database (www.proteinatlas.org)^15,21^.

## A global subcellular map of the metabolic proteome

To generate a global map of the subcellular distribution of the human metabolic proteome with single-cell resolution, we analyzed data that we previously generated for the subcellular section of the Human Protein Atlas (HPA).^15^ This resource contains the subcellular distribution of the human proteome in a wide variety of human cell lines, observed using high-throughput immunofluorescence (IF) and confocal microscopy. Target proteins were detected with the highly validated^22–25^ proteome-wide HPA antibody library.

The Human1 database^26^, a human genome-scale metabolic model, estimates 3,069 genes to be involved in 13,070 biochemical reactions, involving 8,369 metabolites (Supplementary Table 1). Of these genes, we have measured the subcellular distribution for 2,126 proteins (corresponding to 2,126 genes, HPAv23 Subcellular Section, 17,180 IF images, Supplementary Table 2), representing an 8-fold increase compared to other imaging-based subcellular proteomics databases^16,27^ (Fig. 1a, Extended Data Fig. 1a). Additionally it contains twice the number of enzymes compared to recent single-cell MS-based proteomics datasets^28^, which are furthermore unable to also resolve protein subcellular localization^29^. Finally, the number of proteins mapped experimentally to cellular structures is much higher than for UniProt (Extended Data Fig. 1b,c), making the HPA subcellular section an unprecedented resource for studying the heterogeneity of the metabolic protein expression at the single cell level.^15^ To exemplify the type of data and the wide variety of cellular compartments covered, IF images for enzymes from each pathway group are shown in Fig. 1b. The monocarboxylate transporter SLC16A1 can be found in the plasma membrane, while CS, a key enzyme in the citric acid cycle (energy metabolism) and ACAT1 (amino acid metabolism) are detected in the mitochondria. DUT is involved in nucleotide metabolism in the nucleus and CYP51A1 is an enzyme in cholesterol (lipid) metabolism in the endoplasmic reticulum (Fig. 1b). Overall, we found a well-known enrichment of enzymes in compartments of cellular metabolism such as the mitochondria, cytosol and secretory organelles and lower representation of enzymes in the nuclear compartment and cytoskeletal structures (Extended Data Fig. 1d).

**Fig. 1:**
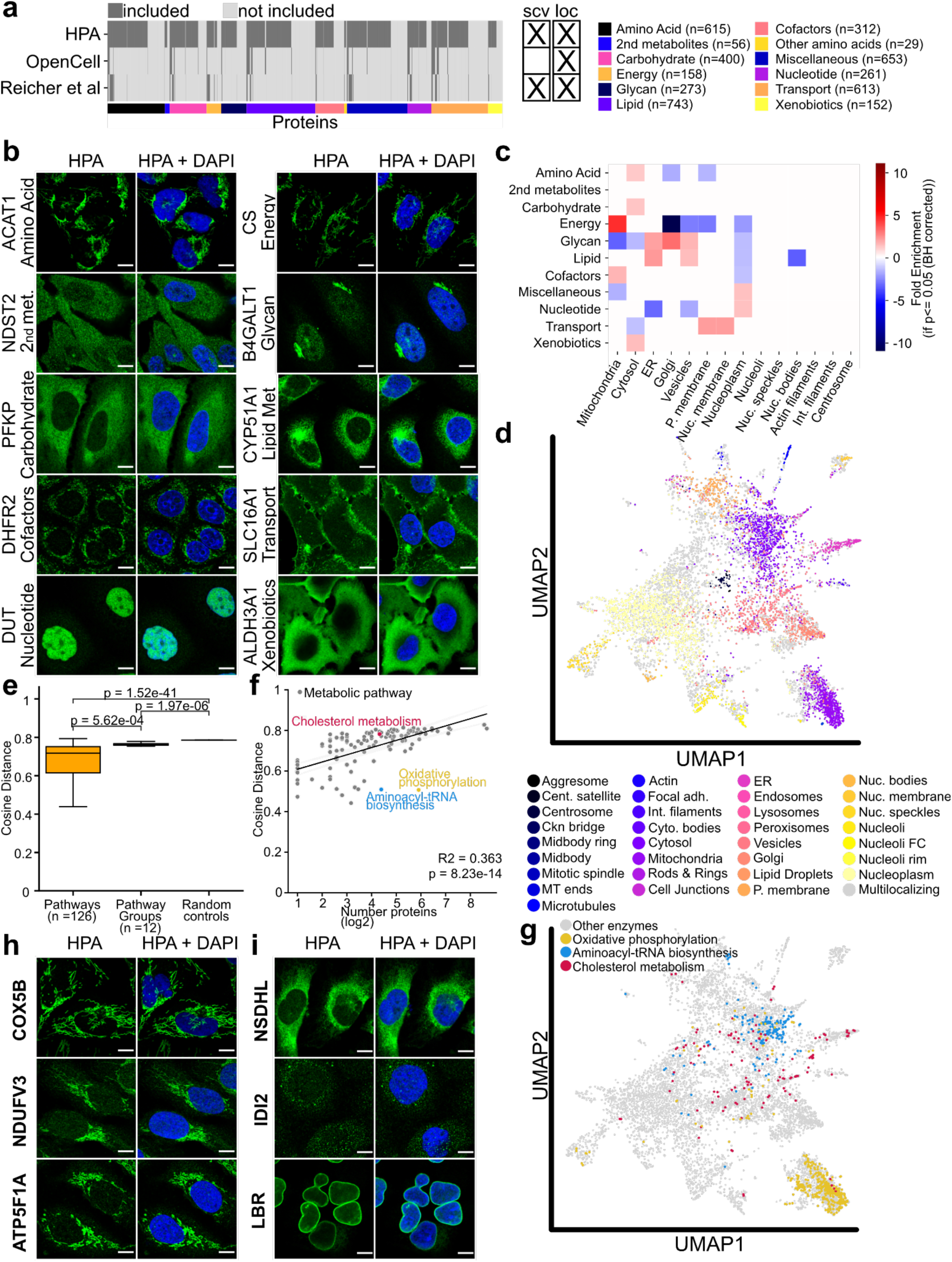
The metabolic proteome. **a,** Coverage of metabolic enzymes across pathway groups in different imaging-based subcellular proteomics dataset. The HPA dataset uses a proteome-wide antibody library and allows for the simultaneous study of single-cell heterogeneity and protein location. The OpenCell^16^ and Reicher^27^ datasets rely on endogenous tagging for protein localization. OpenCell profiles a polyclonal pool of CRISPR/Cas9 edited cells while the Reicher study generated clonal cell populations. As a result, OpenCell is unable to resolve phenotypic cell-to-cell enzymatic heterogeneity from genetic heterogeneity. (scv = single-cell variability; loc = location) **b,** Example images for metabolic enzymes across all metabolic pathway groups, blue = DAPI, green = protein of interest; scale bar corresponds to 10 µm. **c,** Overrepresentation analysis for enzyme localisation in the HPA dataset across different metabolic pathways. Enrichment is shown for significantly overrepresented protein location (P >= 0.05; binomial test). **d,** UMAP visualization of the image features of the entire HPA image dataset.^30^ Metabolic enzymes are visualized. Unilocalizing protein images are colored according to location, while gray data points correspond to multilocalizing proteins. **e,** Pairwise cosine distances calculated for proteins within the same pathway, pathway group, and across random controls in the latent space from image-classification model. Centre line, median; box, first (Q1) and third (Q3) quartiles; whiskers, 1.5× interquartile range (IQR) below Q1 and above Q3. Enzymes within the same pathways are significantly more compartmentalized compared to random protein sets (Mann-Whitney U test; p = 1.52e-41 **f,** Logarithmic correlation between the cosine distance from D and the number of proteins in a pathway. Cholesterol biosynthesis (red), oxidative phosphorylation (blue) and aminoacyl-tRNA biosynthesis (yellow) are highlighted. **g,** UMAP from c, proteins in cholesterol biosynthesis (red), oxidative phosphorylation (blue) and aminoacyl-tRNA biosynthesis (yellow) are highlighted. **h,** Example images for enzymes in oxidative phosphorylation (blue = DAPI, green = protein of interest; scale bar corresponds to 10 µm). **i,** Example images for enzymes in cholesterol biosynthesis (blue = DAPI, green = protein of interest; scale bar corresponds to 10 µm).

## Compartmentalization of the metabolic proteome

A pathway-specific analysis against the background of the metabolic proteome revealed distinct spatial partitioning of metabolic pathways (Fig. 1c). For example, mitochondria harbor many enzymes involved in energy and amino acid metabolism, while transporters are overrepresented in the membrane compartments, where they play a role in shuttling metabolites between different organelles (Supplementary Table 3). To quantify the similarity of subcellular distribution for different sets of metabolic enzymes, we obtained a compressed representation of the spatial information in the entire HPA image dataset from Ouyang et al^30^. We measured the average pairwise cosine distance of these image embedding vectors in the latent space for different proteins (Fig 1d, Extended Data Fig. 1e, Supplementary Table 4). Enzymes within the same metabolic pathway were significantly more similar in their subcellular distribution than random similar-size sets of proteins (Mann-Whitney U; p=1.52e-41; Fig 1e). Therefore, our proteome data confirm metabolic compartmentalization, a well-established biological phenomenon^31^, where similar metabolic reactions are spatially concentrated in similar cellular regions. We identified a correlation between the number of proteins in a pathway and the degree of their compartmentalization (Extended Data Fig. 1f, g; R2 = 0.363, p=8.23e-14; no correlation was observed for random sets of proteins), indicating that pathways with fewer proteins are spatially more constricted whereas pathways with a greater number of proteins have a greater reach across the cell. Interestingly, we identified several outliers from this trend (Extended Data Fig. 1f). For example, oxidative phosphorylation in the mitochondria as well as aminoacyl-tRNA biosynthesis in the cytosol exhibit a significantly elevated level of compartmentalization (Fig. 1f-h). Oxidative phosphorylation is catalyzed by large supramolecular complexes, such as ATP synthase which consists of 17 distinct proteins encoded by 19 different genes.^32^ tRNA charging requires 20 separate enzymes, one for each amino acid. Thus, in both cases, multiple proteins are essentially carrying out a single step in a metabolic pathway. Conversely, cholesterol biosynthesis follows the general trend and is less compartmentalized as it is tightly regulated in the endoplasmic reticulum but occurs in additional organelles such as the peroxisomes or the nuclear membrane (Fig. 1f,g,i).^33^ We have implemented an interactive plugin to explore these embeddings for all metabolic pathways at https://www.proteinatlas.org/humanproteome/subcellular/location+umap.

In conclusion, the subcellular resolution of our dataset allowed us to precisely chart the spatial compartmentalization of the metabolic proteome, revealing that metabolic networks can span several organelles, thereby establishing a complex subcellular landscape. Although partitioning of metabolic reactions has been long known, this is the first time it has been demonstrated at a global metabolic proteome level in single cells.

## Imaging-based subcellular proteomics reveals widespread single-cell metabolic heterogeneity

The single-cell resolution of imaging-based proteomics makes it possible to assess the dynamic properties of the metabolic proteome. Enzymes that are temporally or otherwise regulated in terms of abundance or localization will appear as heterogeneously expressed in our static images of asynchronous cells.

We previously established that a large portion of the human proteome (22.5%, 2,960 of 13,147 proteins) shows significant cell-to-cell variability in protein expression levels or spatial distribution (Fig. 2a, Extended Data Fig. 2a).^15,34^ Strikingly, an even larger percentage of metabolic enzymes across all pathway groups display heterogeneous expression compared to the entire HPA mapped proteome (805 of 2,126 enzymes, 37.9%, *i.e.,* 68.4% higher than whole proteome, p=4.7e-47, binomial test; Fig. 2b). Interestingly, mitochondrial enzymes displayed a particularly high degree of cellular variability (p=7.0e-6, binomial test, BH corrected; Fig. 2c, Extended Data Fig. 2b). To further investigate the biological effects of metabolic heterogeneity on cellular function, we performed a gene ontology (GO)-term enrichment analysis of variably and stably expressed enzymes against the background of the entire metabolic proteome (Supplementary Table 5). For stably expressed enzymes, this analysis revealed a significant enrichment for terms related to protein ubiquitination, DNA damage response and Wnt signaling. For variably expressed ones, we found an enrichment of genes involved in central metabolism (*e.g.,* ATP generation and glycolysis), as well as nucleotide metabolism (Fig. 2d). We employed a random forest regression model to predict protein expression variability based on associated Gene Ontology (GO) terms (Extended Data Fig. 2c). The overall model performance was limited (F1 score = 0.353) thus, the predictive power of GO terms in this context remains inconclusive. Nevertheless, our data suggest a biological role for metabolic heterogeneity, although its origins and exact biological functions remain to be determined.

**Fig. 2:**
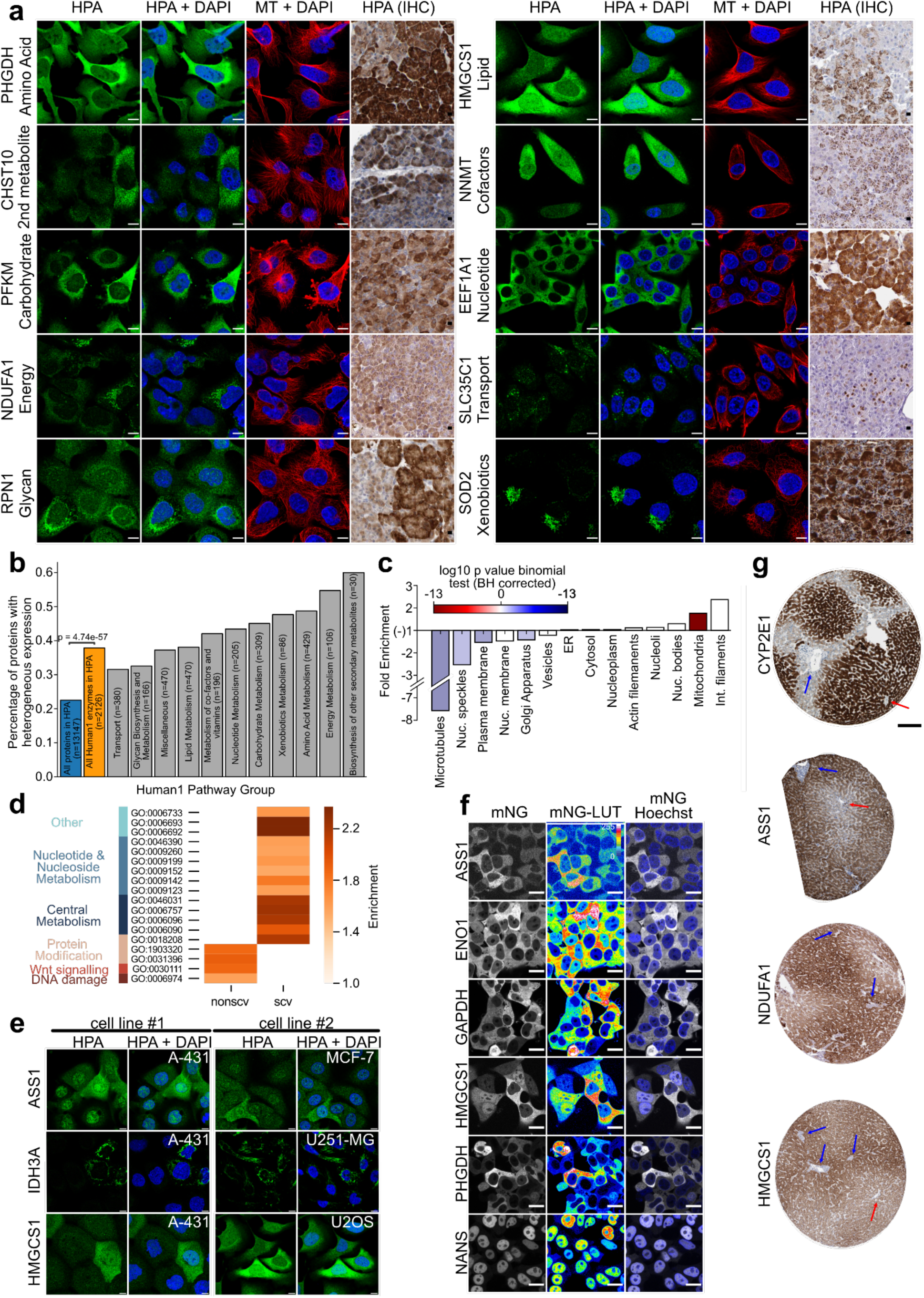
Metabolic heterogeneity establishes phenotypic states. **a,** Example images in cell lines and pancreatic acinar cells for variable metabolic enzymes across all metabolic pathway groups, blue = DAPI, green = protein of interest, red = microtubules; scale bar corresponds to 10 µm. **b,** Enzymes exhibit a higher degree of heterogeneity compared to the human proteome. **c,** Overrepresentation analysis for variable enzyme localisation in the HPA dataset compared to the background of the metabolic proteome. **d,** GO-term enrichment analysis of variably and nonvariably expressed enzymes against the background of the metabolic proteome. Fold-change >1.5 and P <= 0.05 were used as thresholds. **e,** Immunofluorescence images of ASS1 (top), IDH3A (middle) and HMGCS1 (bottom) reveal conserved single-cell heterogeneity across multiple cell lines. **f,** A single cell can reproduce heterogeneous populations, as shown in clonal expansion experiments of mNG-tagged cell lines, grey = mNG tagged protein of interest, blue = Hoechst. **g,** Cellular heterogeneity is conserved in tissue. Immunohistochemistry images for metabolic zonation in liver tissue. Red arrows indicate central vein, blue arrows display portal triads. Scale bar corresponds to 200 µm.

## Metabolic heterogeneity is conserved *in situ* and manifested in the lineage of a single cell

More than half of these enzymes (396 of 733, 54%) showed similar cell-to-cell variation in more than one human cell line (Fig. 2e, Extended Data Fig. 2d), suggesting that they are controlled by conserved regulatory mechanisms. To confirm that the phenotypic heterogeneity is not driven by genetic differences caused by increased mutation rates in cancer cells^35^, we performed a clonal expansion experiment of mNG-tagged Hek293T cells. This key experiment demonstrated that individual cells have the capacity to reproduce highly heterogeneous cell populations, thus establishing myriad functional phenotypes over just a few cell divisions. Heterogeneity observed in the previous dataset was confirmed for several metabolic enzymes belonging to different pathways, including ENO1 and GAPDH (glycolysis), ASS1 (arginine metabolism), PHGDH (serine metabolism), NANS (nucleotide sugar metabolism) and HMGCS1 (cholesterol biosynthesis) (Fig 2f). To further validate metabolic heterogeneity in clonal lineages, we annotated images corresponding to 246 metabolic enzymes from a CRISPR tagging screen in HEK293T and HAP cells by Reicher et al (Supplementary Table 6)^27^. Their results show heterogeneity for over half of the enzymes included in the dataset (53%, 131 of 246). Out of the enzymes showing similar localization to the HPA, a majority show conserved heterogeneity across cell lines (59%, 101 out of 171), similar to our previous analysis across different cell lines in the HPA dataset (Fig 2e). Taken together, these results highlight metabolic heterogeneity as an inherent mechanism of cell biology.

Metabolic heterogeneity *in situ* is impacted by different cell lineages and interactions between multiple cell types. To confirm that human metabolic enzymes exhibit robust phenotypic heterogeneity beyond cell line populations, we analyzed data from a MS-based study of murine liver zonation with single cell resolution.^36^ Here, Rosenberger, *et al.,* identified over 700 proteins with varied expression related to the distance from central vein and portal triads, and over half were enzymes (384 of 742, 51.7%). Out of those enzymes, we observe a significantly higher proportion with variable expression in our dataset (137 of 286, 47.9%, p=0.00034, binomial test) compared to the entire metabolic proteome (805 of 2126 enzymes, 37.9%). For example, the well known zonation markers CYP2E1 and ASS1 display heterogeneity in cell lines (Extended Data Fig. 2e, Fig. 2a,e,f) and increase in expression towards the central vein (CYP2E1) or the portal triad (ASS1) (Fig. 2g). The variable enzymes HMGCS1 (Fig. 2a,e,f) and NDUFA1 (Fig. 2a), an enzyme in oxidative phosphorylation, display similar zonation characteristics to ASS1, with elevated expression towards the portal vein (Fig. 2g). Those findings are in line with previous studies confirming that oxidative phosphorylation is more active in the periportal zone.^37,38^ Furthermore, they exemplify how metabolic enzyme expression heterogeneity can give rise to significant differences in the metabolic capacity of genetically identical cells and thus diversify cellular phenotypes. Altogether, we show that cell line systems are a suitable model to study non-genetic heterogeneity and can serve as a valuable tool to understand cellular plasticity *in situ*.

## Autonomous metabolic heterogeneity is established post-transcriptionally

The cell cycle is a major contributor to cellular heterogeneity, and we interrogated whether it could possibly explain the staggering amount of metabolic heterogeneity in our data using a targeted single-cell proteogenomic screen of the U-2 OS cell line with precise cell cycle mapping over interphase^34^. Strikingly, as much as 76.6% of enzymes (188 of 245) included in the study displayed non-cell cycle dependent (CCD) protein expression heterogeneity. Interestingly, this is the case for so-called “housekeeping genes”, such as enzymes involved in glycolysis or the citric acid cycle, including GAPDH, PDHB and LDHB. They are stably expressed on the transcriptional level, but exhibit significant variability of protein expression^39^ that is independent of cell cycle progression (Fig. 3a). As a result, and in line with previous studies,^12,40,41^ our results show that single-cell transcriptomic data alone is a poor predictor of phenotypic metabolic states. In fact, while the variability in protein expression for non-CCD enzymes is comparable to that for CCD proteins, the variability observed for their corresponding RNAs is lower (p=0.0030, respectively, Kruskal-Wallis test, Fig. 3b) and independent of their overall expression levels (Extended Data Fig. 2f). Furthermore, the physical properties of variable metabolic enzymes differ from CCD proteins, with lower protein disorder fractions and elevated melting temperatures (Extended Data Fig. 2g,h, Supplementary Table 7). Those findings point to fundamental differences in how enzyme variability and protein cycling are regulated, such as through upstream kinase families^42^.

**Fig. 3:**
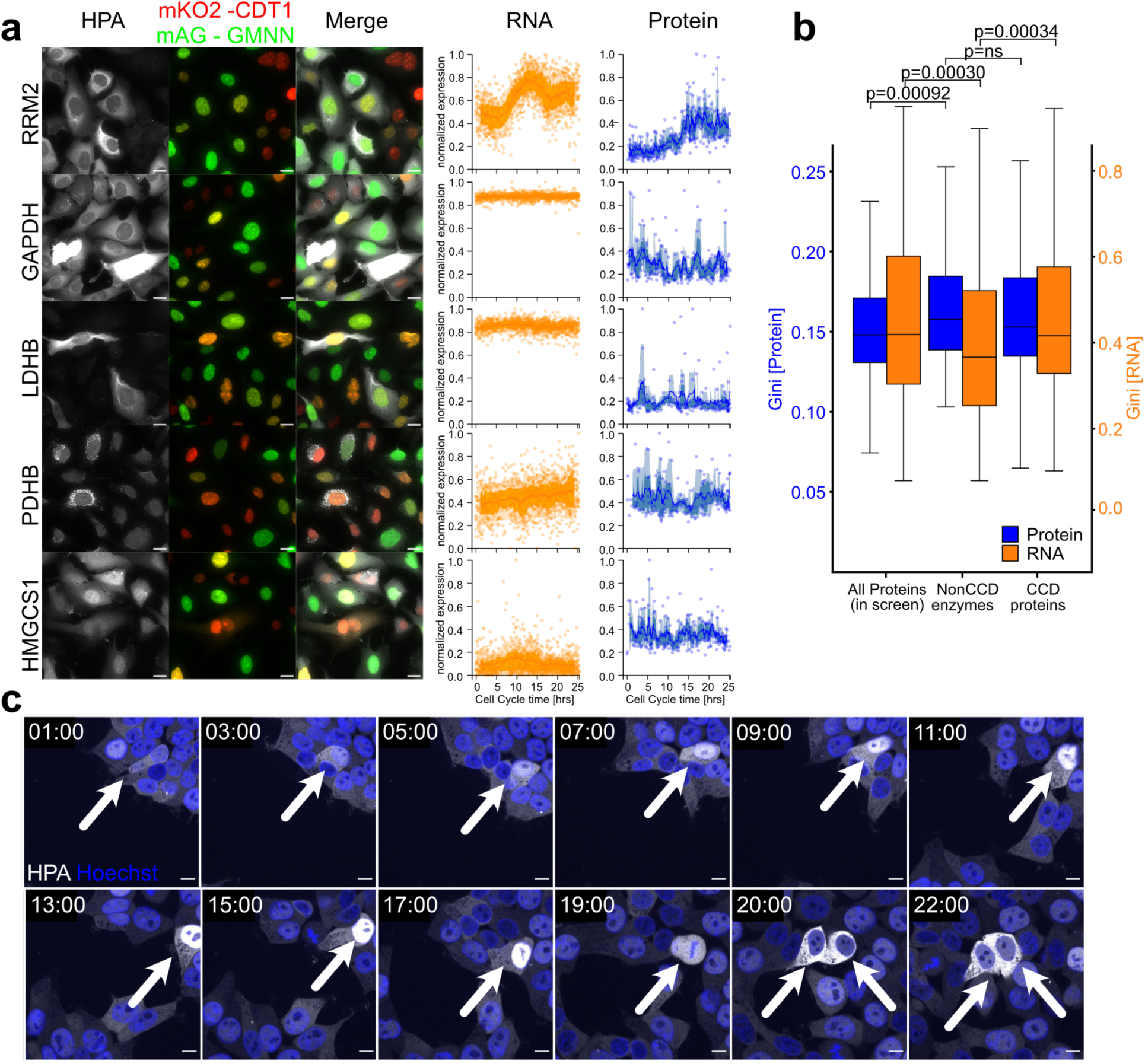
Metabolic heterogeneity is exhibited independently of cell cycle progression and established post-transcriptionally. **a,** Left to middle: Example immunofluorescence images for CCD (RRM2) and non-CCD enzymes (GAPDH, ENO1, PDHB, HMGCS1) in U2OS-FUCCI cells from Mahdessian *et al*^34^, green = mAG-GMNN, red = mKO2-CDT2, grey = protein of interest; scale bar corresponds to 10 µm. Middle and right, single-cell temporal expression profiles for RNA (orange) and protein (blue). Line, moving mean; darker shade, 25th to 75th percentile range; lighter shade, 10th to 90th percentile range; points, individual cell data. **b,** Transcript profiles (orange) do not reflect the variability observed at the protein level (blue), and in fact while variability of non-CCD enzymes is higher than all proteins, the variability of their corresponding RNAs is lower than all proteins in the screen. For box plots: center line, median; box, first (Q1) and third (Q3) quartiles; whiskers, 1.5× interquartile range (IQR) below Q1 and above Q3 **c,** Time-lapse imaging data for mNG-tagged HMGCS1 protein (white) demonstrates cytoplasmic localization in G1, and then translocation to the nucleus (blue); scale bar corresponds to 10 µm. However, the overall expression levels of HMGCS1 remain stable, corroborating the findings of cell cycle independent expression heterogeneity from a.

We investigated differences in PTM regulation of metabolic enzymes compared to cycling proteins by comparing the kinases upstream of phosphosites on these proteins (Supplementary Table 8). While cycling proteins are enriched for CMGC kinases upstream of phosphosites (p=5.0e-15), which was reported previously^34^ and is tied in function to regulating cell fate^43^, metabolic enzymes are enriched for AGC and TK kinase families (p=0.0016 and 0.00078), which have respective functions in metabolism^44^ and cell signaling^45,46^. Enzymes that exhibit variability between individual cells display a strong enrichment for upstream TKs when compared to non-variable enzymes (p=6.9e-7). This indicates that metabolic enzymes have distinct PTM regulation and that the regulation of enzymes displaying variable expression between individual cells is regulated by machinery involved in cell signaling transduction and cell-to-cell communication.

Importantly, the cell cycle and metabolic heterogeneity can still influence each other. HMGCS1 is an enzyme involved in cholesterol biosynthesis that has variable, non-CCD expression levels of both RNA and protein (Fig. 3a). However, time lapse microscopy reveals a coordinated CCD translocation of cytoplasmic HMGCS1 to the nucleus during interphase (Fig. 3c), showing how different layers of heterogeneity can come together in the same cell. In this case, the differences in expression levels establish differences in metabolic capacity between individual cells, while the location of the protein is linked to cell cycle position. This result also highlights the fact that gene expression heterogeneity is neither able to capture the subcellular localization of a protein nor the spatial heterogeneity established by protein translocation. In conclusion, we find that metabolic heterogeneity is largely established independently of the cell cycle, mostly regulated post-transcriptionally, for example through translocation^47^, targeted degradation^48^, and post-translational modification^48^ of enzymes.

## Non-genetic enzymatic heterogeneity establishes continuous metabolic states in cell populations

We observe cellular heterogeneity across all metabolic pathways which could give rise to a plethora of theoretical metabolic states. To investigate these, we performed 4i multiplexed imaging and stained for rate limiting enzymes across 3 metabolic pathways: SOD2 (ROS detoxification), ACACA (fatty acid synthesis) and PCK2 (gluconeogenesis), along with tubulin and DAPI (Extended Data Fig. 3a,b). For each condition, we profiled three additional metabolic proteins, 135 in total (Fig. 4a, Supplementary Table 9). We extracted single cell morphology and staining features for all individual cells (n=46,274) and embedded them in a two-dimensional feature space (Fig. 4b, c). There were no batch effects and high reproducibility across replicates (mean pairwise Pearson correlation between the median features extracted from all replicates = 0.995; Extended Data Fig. 4a,b).

**Fig. 4:**
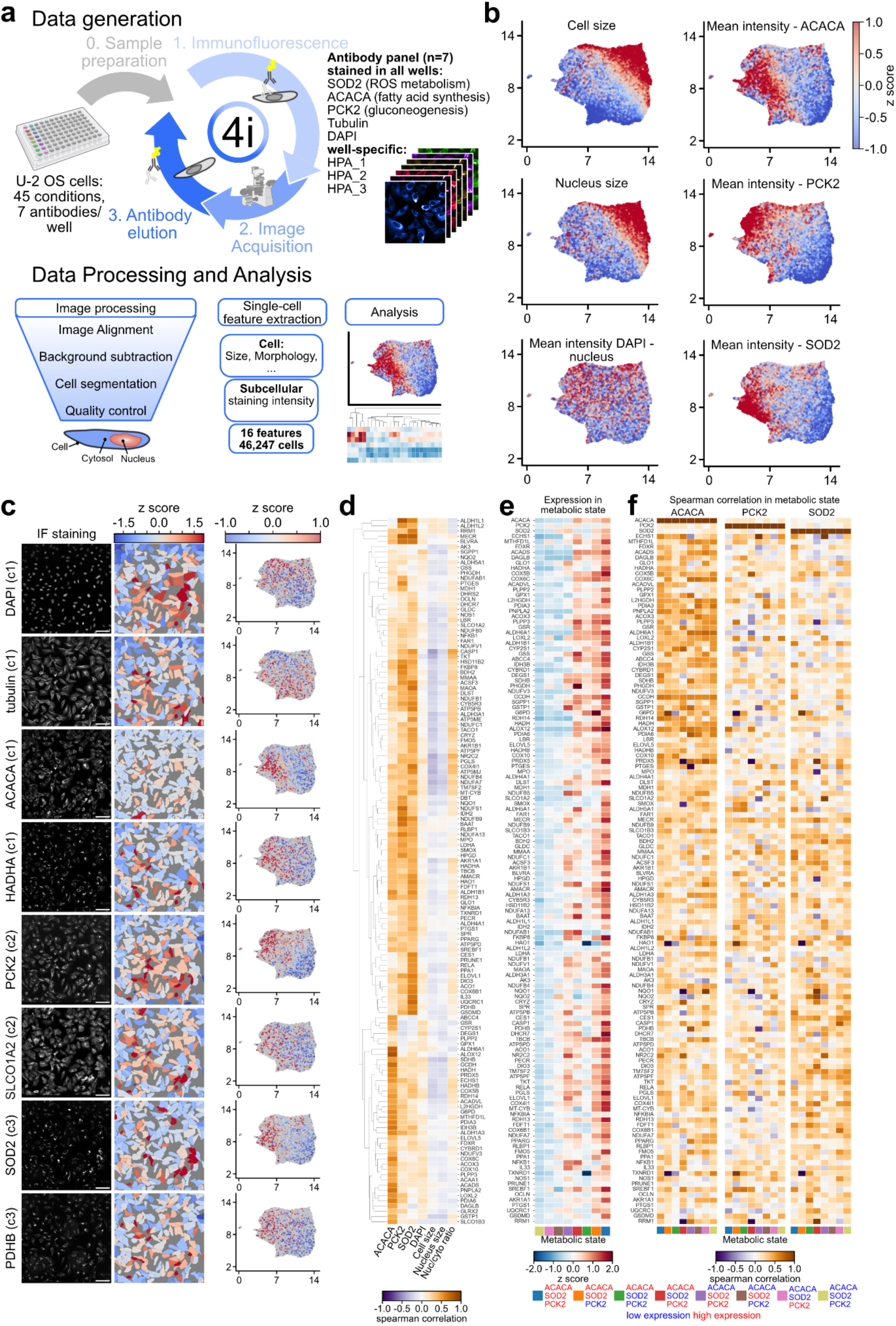
Metabolic heterogeneity establishes a continuous metabolic state landscape. **a,** Experimental Design of a three cycle 4i experiment. Wells were stained for three rate limiting enzymes (SOD2, ACACA and PCK2), and three well-specific additional enzymes, along with DAPI, and tubulin. Cells were segmented and single-cell morphological and staining features were extracted for analysis. **b**, cells were projected into a 2D feature landscape using UMAP dimensionality reduction on single cell features. UMAP data points colored by different features. **c**, Example data from one well; left: immunofluorescence images for 7 antibodies and DAPI from one 4i experiment (greyscale, scalebar corresponds to 30 µm); middle: mean intensity per cell visualized on the cell masks from the left panel; right: mean intensity per cell visualized on UMAP obtained in b). **d**, Heatmap visualizing spearman correlation between mean protein and metabolic marker expression. **e**, Heatmap displaying the mean expression (z score) of proteins across 8 metabolic clusters defined by high/low expression of the three metabolic markers. **f**, Heatmap displaying the spearman correlation between protein and metabolic markers (from left to right: ACACA, PCK2 and SOD2) in each metabolic cluster.

The expression of the selected metabolic markers is not mutually exclusive across individual cells, suggesting that cellular metabolism operates along a continuum, or that a more comprehensive panel of markers is required to fully capture the range of possible states. To probe the relationships between these pathways, we calculated pairwise correlations between enzyme expression and other cellular features in each separate 4i experiment. Metabolic enzyme levels were only loosely associated with cell or nuclear size (Fig. 4d), in line with previous studies describing decreased metabolic activity at larger cell sizes^49^. However, the expression of a subset of enzymes, including several enzymes from the TCA cycle and oxidative phosphorylation correlate positively with the cytoplasmic-to-nuclear aspect ratio.

The enzymes included in our screen exhibit distinct relationships to the markers used in this study (Fig. 4c, d) suggesting that different metabolic pathways are activated in single cells and that metabolic states are not defined by active or inactive metabolism. For example, PDHB correlates positively with SOD2 (ROS metabolism) and PCK2 (gluconeogenesis) levels but its expression is independent of ACACA (fatty acid synthesis) (Fig. 4c,d). Similarly, the expression of cytosolic antioxidant enzymes such as GSR, GSTP1, and GLO1 is independent of the mitochondrial antioxidant enzyme SOD2, but correlates strongly with ACACA, a rate limiting enzyme in fatty acid synthesis (Fig. 4d). This suggests a decoupling of mitochondrial and cytosolic ROS metabolism due to metabolic compartmentalization. Elevated fatty acid synthesis in cells with high ACACA levels leads to increased oxidative stress and thus requires upregulating cytoplasmic ROS metabolism to preserve the redox balance^50^.

To further investigate whether proteins can perform metabolic state specific functions, we separated the cells into 8 metabolic states based on the relative expression of SOD2, PCK2 and ACACA (high expression corresponding to positive z-scores, low expression to negative z-scores) (Extended Data Fig. 4c), and characterized the expression of all measured enzymes across these states. Overall, the state with low expression of all three markers exhibited the lowest global enzyme activity (Fig. 4e), suggesting the existence of an “inactive” state. Higher expression of one or more markers gave rise to distinct active states with subtle variations in enzyme expression and the relationship between different metabolic states across states, implying that the activity of individual metabolic pathways is context-dependent (Fig. 4f). For example, while PDHB generally correlated with SOD2 and PCK2 across the population, its expression was negatively associated with PCK2 in a specific state (high SOD2, high PCK2, low ACACA).

Taken together, this multiplexed map of subcellular metabolism provides a comprehensive view of the metabolic state landscape at the single-cell level. Future studies are required to dissect the functional roles of these states, for example by studying their dynamics across a variety of different drug perturbations. Nevertheless, this widespread variation in enzyme expression, and the state specific correlation between different metabolic pathways create a need to reinspect the traditional static wiring of metabolic flux diagrams and may change our understanding of the metabolic regulation of cell types and single cells (Extended Data Fig. 5a, b).

## The intricate subcellular landscape of the metabolic proteome

Multilocalizing enzymes, including HMGCS1, open up intriguing questions about the interconnectedness of enzyme subcellular distributions and corresponding metabolic processes. With the subcellular resolution of imaging proteomics, we found that more than half of all metabolic proteins are multi localizing (53.8%, 1,144 of 2,126; Fig. 5a,b), which is comparable to the entire proteome (56.5%, 7,434 of 13,147, p=0.011, binomial test; Extended Data Fig. 6a)^15,16,51,52^. Many enzymes localize to both the cytoplasmic and nuclear compartments, such as ALAS1 (mitochondria and nucleoplasm), NDUFB4 (mitochondria and nuclear membrane) and GCH1 (cytosol and nucleoplasm), while other enzymes localize to multiple structures in one meta-compartment, such as the transporter SLC7A5 (cytoplasm and vesicles; Fig. 5a). As mentioned previously, enzymes are overrepresented in the cytoplasm (p = 5.0e-31, Fisher exact) and are less likely to be present in the nucleus compared to the entire human proteome (p = 1.3e-28, Fisher exact; Extended Data Fig. 1d). However, multilocalizing enzymes are more likely to be present in the nucleus compared to unilocalizing enzymes, *i.e.,* those found in a single location (Extended Data Fig. 6b).

**Fig. 5:**
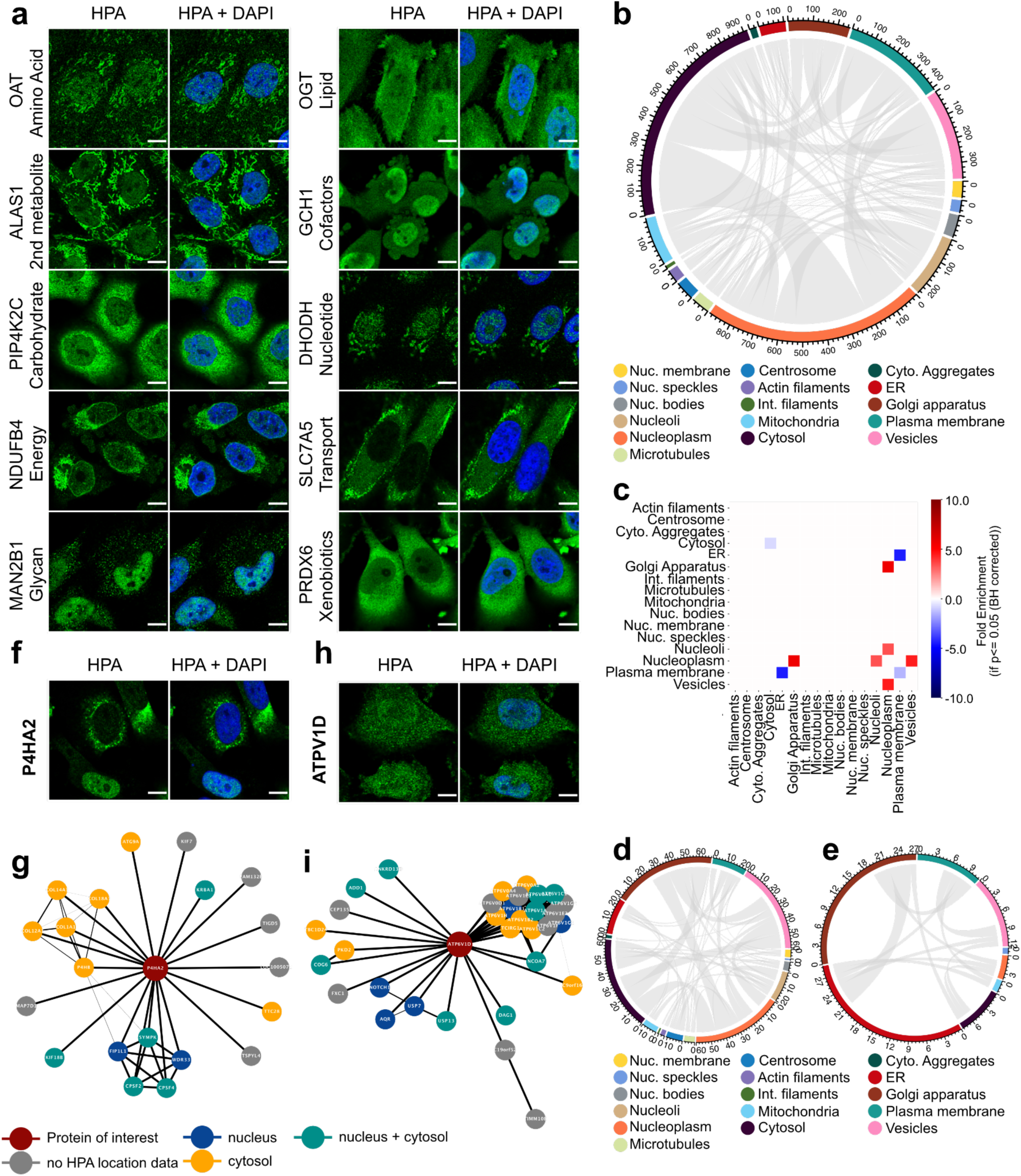
The intricate subcellular landscape of the metabolic proteome. **a,** Example images for multilocalizing metabolic enzymes across all metabolic pathway groups (blue = DAPI, green = protein of interest; scale bar corresponds to 10 µm). **b,** Circos plots with the identified proteins of each compartment presented and sorted by meta-compartments (cytoplasm = red, secretory pathway = green, nucleus = blue). Multilocalizing proteins are connected by a line (secondary colors represent multilocalization across metacompartments). The plots show connections among metabolic proteins in the HPA dataset. **c,** Overrepresentation analysis for enzyme co-localisation in the HPA dataset compared to UniProt localisation information. Enrichment is shown for significantly overrepresented location pairs (P >= 0.05; binomial test, BH corrected). **d, e,** Circos plots with the identified proteins of each compartment presented and sorted by meta-compartments (cytoplasm = red, secretory pathway = green, nucleus = blue). Multilocalizing proteins are connected by a line (secondary colors represent multilocalization across metacompartments). The plots show connections among **(d)** enzymes involved in glycan biosynthesis in the HPA dataset and **(e)** enzymes involved in glycan biosynthesis with experimental evidence on UniProt. **f,** P4HA2 localizes to the ER and the nucleus (blue = DAPI, green = protein of interest, scale bar corresponds to 10 µm). **g,** Protein-protein interaction network of interactors of P4HA2, colored by protein localization. P4HA2 interacts with a cluster of proteins in the cytoplasm and the nucleus. **h,** ATP6V1D localizes to the cytosol and the nucleus (blue = DAPI, green = protein of interest, scale bar corresponds to 10 µm). **i,** Protein-protein interaction network of ATP6V1D, colored by protein localization. P4HA2 interacts with a cluster of proteins in the cytoplasm and the nucleus.

## Multilocalizing enzymes are functionally understudied

Our data indicate that 54% of proteins localize to multiple compartments, which is likely an underestimation, since spatiotemporal protein expression varies across cell types and contexts^53^. We define multilocalizing proteins as potentially multifunctional proteins. Text-mining studies predict that over a quarter of all proteins can carry out multiple functions^54,55^ establishing a great discrepancy between the number of multilocalizing proteins and the number of multifunctional proteins with a strong overrepresentation of enzymes among known moonlighting proteins (39 of 86, 45%, p = 2.1e-10, binomial test)^56,57^. We compared the location data from the HPA subcellular section to experimentally verified location data extracted from the curated database UniProt (after subtracting locations derived from HPA; Supplementary Table 10). We found a significantly higher proportion of multilocalizing proteins in the HPA dataset than in UniProt across all metabolic pathways (p = 9.8e-32, Fisher exact, Benjamini-Hochberg (BH) corrected; Extended Data Fig. 3c) with an overrepresentation of nuclear localizations (Fig. 5c) across many metabolic pathways (Fig. 5d,e, Extended Data Fig. 7a-i). For example, IF imaging revealed a localization to both the cytosol and nucleus for ALDH3A1 and ACLY, as well as to the Golgi apparatus and nucleus for CERT1 validated with multiple independent antibodies, while there is currently no experimental evidence for a nuclear location in the UniProt database for any of those proteins (Extended Data Fig. 6d). Those findings suggest unknown nuclear metabolic activity for many enzymes, as recently reported for TCA cycle enzymes^47,58^. We confirmed these findings in our data and show that 29% (12 of 41) TCA cycle enzymes localize to both the mitochondria or cytosol and the nucleus (Extended Data Fig. 8a,b). The large amount of protein multilocalization could also point to uncharacterized non-metabolic nuclear functions for metabolic enzymes,^59^ similar to the recently described involvement of Golgi proteins in DNA damage^60^. Interestingly, multilocalizing enzymes are more likely to exhibit variable expression than enzymes localizing to a single cellular compartment (35.8.% of unilocalizing proteins are variably expressed, 39.8% of multilocalizing; p = 7e-4 Fisher’s exact test). The combination of cellular heterogeneity and multilocalization can greatly expand the functional diversity of cells in a population.

The variable expression patterns of ENO1, GAPDH and PDHB in the nucleus and cytoplasm (Extended Data Fig. 8c) serve as great examples of how the spatiotemporal regulation of enzymes establishes diverse functional capacities among cells. PDH is a complex that generates acetyl-CoA in the mitochondria and the nucleus. Mitochondrial acetyl-CoA is used in the TCA cycle while nuclear acetyl-CoA is crucial for histone acetylation^61^. Our data show stable expression of PDH subunit beta (PHDB) in the nucleus, while mitochondrial PDHB levels vary between individual cells. These findings indicate a decoupling of the two processes (Extended Data Fig. 8c) and compartment-specific regulation of PDHB levels. In other cases, multilocalizing proteins can perform distinct functions in different compartments. The glycolytic enzyme GAPDH is one such multifunctional enzyme; it carries out its canonical glycolytic function in the cytosol, while it is also involved in apoptotic pathways and transcriptional gene regulation in the nucleus (Extended Data Fig. 8c)^62^. ENO1 is another glycolytic enzyme in the cytosol, but it also functions as a plasminogen receptor on the cell surface and has RNA/DNA regulatory functions in the nucleus^63^. Altogether, our analysis reveals new insights into multilocalizing characteristics of metabolic enzymes and highlights the need for additional studies to decipher their non-canonical functions.

## Identification of multifunctional enzymes

Multifunctional enzymes perform multiple functions in different parts of the cell^64^. To find such proteins, we combined localization information from our global spatial proteomic data with protein-protein interaction (PPI) measurements that capture the molecular-scale local environment of proteins. We assembled enzyme interaction partners from various studies^16,65,66^, as well as from the STRING database^67^ (filtered for physical interactions; experimental evidence and confidence > 0.7), and constructed interaction networks for each multilocalizing metabolic enzyme (Supplementary Table 11). This allowed us to assess non-metabolic functions using a guilty-by-association approach. For example, P4HA2, an enzyme involved in collagen synthesis in the ER (Fig. 5f), interacts with other members of the collagen synthesis pathway in the cytoplasm. However, in the nucleus, it interacts with members of the CPSF complex and may be involved in mRNA polyadenylation as well as alternative splicing (Fig 5g, Extended Data Fig. 8d). Interestingly, recent studies have revealed that many metabolic proteins are capable of RNA binding, indicating that a large number of enzymes could have non-canonical functions in cell biology.^68,69^ Another example is ATP6V1D, which is a subunit of the vacuolar ATPase, a proton pump responsible for controlling the intra- and extracellular pH of cells (Fig. 5h). It interacts with NOTCH1 and USB7 in the nucleus and could thus play an additional role in the regulation of NOTCH signaling (Fig. 5i, Extended Data Fig. 8e). Altogether, combining spatial proteomic and interactomic analyses revealed potential non-canonical protein functions, including for well characterized enzymes such as P4HA2 and ATP6V1D, and provides a promising avenue for future systematic studies of multifunctional enzymes.

## Exploration of metabolic proteome phenotypes in the HPA database

This image-based map of the metabolic proteome is freely available as part of the Human Protein Atlas database (www.proteinatlas.org/humanproteome/subcellular/metabolic+proteome v24). The subtle variations in protein expression and subcellular localization between cells from undisturbed, well-characterized cell lines can hint at novel protein functions and cell-specific regulatory mechanisms. These findings offer a robust baseline for future research into genetic, environmental, and pharmacological perturbations of metabolic cellular phenotypes. Moreover, as antibody-based staining of histological samples is a routine procedure in clinical contexts, our resource allows for direct comparison between cell lines and tissues. Altogether, these data will facilitate constructing spatially resolved proteome-wide metabolic flux models to better understand heterogeneous drug responses in cell populations and ultimately provide new opportunities to drug specific cell states.

## Conclusion

In this study, we describe the heterogeneity of the human metabolic proteome on single-cell and subcellular levels. We show pervasive non-genetic heterogeneity of metabolic enzyme expression. Over half of all enzymes localize to multiple locations in human cells, including several newly identified multifunctional proteins, and nearly two fifths exhibit cell-to-cell variable expression that is largely autonomous of the cell cycle. This heterogeneity is recapitulated in many cell lines, native tissue contexts, and clonal expansions of individual cells, appears to be established at a post-transcriptional level, and gives rise to continuous metabolic states. This imaging-based subcellular proteomic dataset will be a valuable resource across many areas of research, including cell biology, metabolic modeling or pathology.

The compartmentalization of metabolism is a well established biological concept, yet global maps of metabolic compartmentalization are still lacking. In this study, we provide experimental evidence for the subcellular localization of over 2,100 metabolic enzymes, demonstrating global metabolic compartmentalization on the proteomic level. While the enzyme activity is certainly linked to subcellular localization and colocalization with other enzymes, it also depends on the presence of respective precursors, products and inhibitors. Recent developments in MALDI-MSI allow for the subcellular profiling of lipids and other metabolites with single cell resolution^2,13^, but no database for the subcellular localization of metabolites exists. Subcellular fractionation followed by lipidomic and proteomic analysis revealed that more than half of the proteins and almost all lipids localized to multiple cellular compartments^70^, probably due to easier diffusion within the cell. Future work is required to dissect the role of metabolite compartmentalization^71^. Additionally, enzyme activity can be regulated by post-transcriptional modifications, such as phosphorylation, which remains insufficiently characterized^42^, especially with regards to spatial information. We showed that enzymes displaying cell-to-cell variable expression are particularly enriched for specific upstream tyrosine kinases involved in cell signaling when compared to all enzymes.

Protein localization is an important layer of cellular regulation that cannot be inferred from transcript or protein expression data alone^14^. We find that over half of all metabolic enzymes localize to multiple compartments, greatly expanding their context-dependent functionalities. Even so, the secondary functions of enzymes are rarely described. We show that compartment-resolved PPI data have the potential to unravel novel functions for multilocalizing proteins and provide a suitable starting point for in-depth functional studies of protein multifunctionality. Importantly, our imaging-based spatial proteomics approach does not distinguish between proteoforms^72^. As such, protein multilocalization and multifunctionality may be driven by post-translational modifications^73^ or structural changes.^74^

Cellular metabolism is an extremely dynamic process that can change over the cell cycle, circadian rhythm, or following signaling events, yet global maps of metabolic variability are missing. In this study, we provide experimental evidence that 805 metabolic enzymes demonstrate variability in expression in terms of abundance or localization. We show that the heterogeneity in enzyme expression establishes continuous metabolic states with state-specific relationships between metabolic pathways. This widespread variation in enzyme expression creates a need to reinspect the wiring of metabolic flux diagrams (*e.g.,* for pathways illustrated in Extended Data Fig. 3 and 5). Metabolic flux is often assessed using bulk transcriptomic data, which is unable to resolve cell-to-cell enzymatic heterogeneity^75^. However, single-cell transcriptomic data alone is unlikely to accurately capture metabolic states, since it is a poor predictor of single-cell protein expression^34,76–79^ and metabolite levels^12^. Consequently, our single-cell proteome-wide metabolic map, although not obtained from the same individual cells, offers an important approximation of cellular metabolic heterogeneity and provides a valuable step towards improved metabolic modeling^75^ which may change our understanding of the metabolic regulation of cell types.

Inherent heterogeneity of metabolic protein expression may serve the purpose of a bet-hedging evolutionary strategy that allows entire cell populations to survive in rapidly changing environments, such as during drug treatments^80,81^, as has been demonstrated in yeast,^82^ bacteria^83^ and cancer cells.^84^ Sufficient heterogeneity may contribute to the resilience of the cell population as a whole. Reduced heterogeneity (*e.g.,* in aging, stress, or disease) may deplete the potential of the cell population to restore cell function and survive under a wider range of conditions. At the same time, the excessive heterogeneity (e.g. observed in aging yeast cells^85,86^ might indicate that the cells are no longer able to maintain the tight regulation of their metabolism that is necessary for effective and efficient cell functioning.

An immense number of possible metabolic states are possible, given the 800 metabolic enzymes that we show to have variable expression in separate experiments and the continuum of metabolic states that we observed using co-stainings for up to 6 enzymes using 4i. Even so, established biochemical frameworks describe a relatively small number of high-level, often mutually exclusive metabolic states (switches), such as glycolysis-dominated or respiration-dominated energy production. Co-regulated enzymatic activity within a metabolic pathway can lead to co-variation of enzyme expression. In this case, it may be possible to pinpoint highly variable metabolic enzymes that are markers for distinct high-level metabolic states within the cell population. However, this co-variation is not absolute; one enzyme can show relatively higher expression than other enzymes in the same pathway if its product is required for another metabolic pathway or if it is performing several different functions^87^. Mapping such variations in the same single cells using multiplexed imaging strategies as proposed in this study, or single cell mass spectrometry-based technologies^88,89^ may reveal the complex landscape of more granular metabolic phenotypes, including the connections, dependencies and compensatory interactions between different metabolic pathways. These diverse enzymatic expression patterns may also give rise to convergent metabolic phenotypes. This phenomenon was recently described for the protein components of the circadian clock, each of which can have widely varying turnover rates in individual cells yet show robust circadian rhythm periods as a whole population.^90^ Converged high-level phenotypes like the circadian rhythm are still underpinned by diverse expression of their individual components, which notably opens up diverse response potential for cells when those individual components are targeted by drug treatments. Better understanding cellular metabolic states facilitates manipulating them and developing novel treatment strategies for diseases, such as cancer, to improve the efficacy of existing drugs and overcome drug resistance.

Measuring metabolic states is a difficult task, and the ways in which metabolic states can converge based on variable enzyme expression, such as was described above for circadian rhythms, point to the potential of enzyme (co)expression and (co)localization to be useful proxies for studying metabolic states. Additionally, the simultaneous spatial profiling of multiple omics layers may be able to measure these proxies and provide novel insights into the regulation of cellular metabolism.^91^

In conclusion, the single-cell map of autonomous enzymatic variability presented in this paper underlines the importance of studying metabolic heterogeneity in many areas of cell biology. This new perspective could revolutionize many areas of medicine, including the development of new treatment designs to avoid drug resistance.

## Materials and Methods

### HPA antibodies and validation

All antibodies within the HPA project are rabbit polyclonal antibodies derived after immunization with recombinant protein epitope signature tags (PrEST) as antigens and purification using the antigen as an affinity ligand.^22^ Antibodies were then quality-controlled for sensitivity and crossreactivity using western blotting and protein arrays before used in immunocytochemistry experiments for the Subcellular Section of the Atlas. The subcellular protein localization results are assigned a gene-specific confidence score. If there were no independent experimental data on UniProt contradicting the results obtained in the immunofluorescence experiment, the antibodies were labeled “approved”, if the independent supported the results, the antibodies were labeled “supported”. Antibodies validated according to the strategies outlined by the International Working Group for Antibody Validation (IWGAV)^92^ were labeled “enhanced”. These strategies include i) validation by co-staining with fluorescently tagged protein; ii) validation by gene silencing or knock out; iii) validation by immunocapture and mass spectrometry and iv) validation by independent antibody.^92^

### Subcellular localization of proteins using immunofluorescence

The distinct localization patterns for each protein assigned on the basis of annotations of the target protein in the HPA Subcellular Section (Supplementary Table 2). In short, a standardized protocol for immunofluorescence was applied^93^ and fluorescent images with confocal resolution were acquired as described in Thul et al^15^. Finally, the protein localizations were manually determined based on the signal pattern of the protein of interest in relation to cellular markers (DAPI, microtubules and the ER) and the negative control following strict annotation guidelines, as previously described^15,94,95^.

### Identification of proteins with cell-to-cell heterogeneity as well as multilocalizing proteins

Protein cell-to-cell heterogeneity was manually identified in images from the HPA Cell Atlas (HPA v23) following strict annotation guidelines as previously described^15,34,95^. In short, cell-to-cell heterogeneity was identified either as variation in staining intensity (= protein abundance) or variation in subcellular distribution between cells in the same microscopic image. Multilocalizing proteins were defined as proteins being present in multiple locations within or across all cell lines and/or images in the Subcellular Section of the HPA. In the context of this study, “heterogeneity” and “multilocalization” thus refer to distinct protein properties and are treated as categorical variables.

### Quantification of protein expression levels in single cells

We quantified the expression heterogeneity in single U-2 OS cells and nuclei in the HPA dataset to validate the manual annotations of single-cell heterogeneity. First, we extracted single-cell and nuclear masks from the HPA images using a previously described DPN-UNet segmentation model^96^, available at https://github.com/cellProfiling/hpa-cell-segmentation. We segmented 23159 images corresponding to 9752 proteins stained in the U-2 OS cell line, resulting in 256969 single cells after filtering out border-touching cells. To assess single-cell heterogeneity, we computed the median absolute deviation (MAD) across all cells in each image. Finally, we compared the MAD between stably and variably expressed enzymes or proteins. The compartment used for the analysis was based on the manual HPA annotations for each protein.

### Simplifying the HPA protein location data

We extracted the protein location data from HPA v23 and collapsed the annotations for all 35 subcellular structures into 14 major organelles (Mitochondria, Cytosol, Cytoplasmic Aggregates, Endoplasmic Reticulum (ER), Golgi Apparatus, Vesicles, Plasma membrane, nuclear membrane, Nucleoplasm, Nucleoli, Nuclear speckles, nuclear bodies, Actin filaments, Intermediate filaments, Centrosome) that were used for the analysis according to Extended Data Table 1.

### Definition of the metabolic proteome, pathways and pathway groups

The metabolic proteome was defined according to the Human1 database, a human genome-scale metabolic model containing 13,070 biochemical reactions with 8,369 metabolites and over 3,069 genes.^26^ The Human1 database provides evidence about the metabolic reactions and pathways for each enzyme and thus allows for grouping the enzymes into different metabolic pathways and pathway groups (Supplementary Table 1). 2,125 genes have corresponding protein localization data in the HPA Subcellular Section v23 (Fig. 1a). The good coverage for enzymes across all pathway groups in the HPA Subcellular Section v23 enables a detailed analysis of the metabolic proteome using imaging-based Spatial Proteomics.

### Subcellular location overrepresentation analysis

For the location overrepresentation analysis we extracted the number of proteins for each subcellular location across our test and background datasets. We calculated the overrepresentation of the location in the test vs the background protein set and performed a two-sided binomial test to test for statistical significance. The assumptions of this test were satisfied as follows: the data were large and nominal, the sample size was less than the population size, the samples were independent, and the probability of a given outcome did not affect the probability of the other. A p-value < 0.05 after Benjamini Hochberg correction was used as a cutoff. The significant enrichment for all locations in the test set versus the background set were visualized in a heatmap or barplot.

### Cosine distance analysis

Ouyang et al embedded the spatial information from the entire HPA image dataset into a compressed representation using a machine learning approach^30^. We obtained these embeddings and visualized them as a 2D projection of a UMAP (Extended Data Fig. 1e). We computed the average pairwise cosine distance (1 - cosine similarity) of all images belonging to genes from the same metabolic pathway or pathway group (Supplementary Table 4). We performed a random control by picking the equivalent number of random images and computing the average cosine distance across 1000 permutation experiments. Analysis was performed with *numpy* and *cupy* for GPU acceleration. We also implemented an interactive UMAP plugin to explore these embeddings across all metabolic pathways (https://www.proteinatlas.org/humanproteome/subcellular/location+umap).

### GO term enrichment and association analysis

We performed a GO term enrichment analysis for biological processes using GOrilla^97^ and filtered for terms with a fold enrichment >1.5 and a p value < 0.05 after Bonferroni correction (Supplementary Table 5). The significant terms were manually grouped into categories (*e.g.,* cell cycle, metabolism) and visualized in a heatmap.

To assess the predictive power of GO term annotations on protein expression variability, we performed a supervised classification using a random forest model. GO term annotations were retrieved from BioMart and filtered for “Biological Process” terms. For each gene, associated GO terms were concatenated into a single space-separated string and vectorized using CountVectorizer (binary=True), encoding presence/absence of terms. The Human Protein Atlas annotations of single-cell variability were binarized into a target variable (1 = variable, 0 = stable). A random forest classifier (n_estimators=200, random_state=42, class_weight=None) was trained and evaluated using 5-fold cross-validation with F1 score as the performance metric (due to class imbalance). The model was implemented using scikit-learn v1.5.2. Following training, GO term feature importances were extracted from the fitted model and ranked by mean decrease in impurity to identify terms most predictive of protein expression heterogeneity.

### Comparison of metabolic heterogeneity with cell cycle heterogeneity

We obtained a list of cell cycle dependent (CCD) and independent (non-CCD) variable proteins^34^. By extracting the single-cell protein and RNA measurements from this study, we were able to compare the degree of variability on the proteomic and transcriptomic level across different non-exclusive groups of human proteins as defined by our data (all proteins in the HPV v23, enzymes in the HPA, enzymes exhibiting cell-cell heterogeneity in HPA v23, and CCD proteins) using Kruskal-Wallis. Furthermore, we obtained the physical properties of human proteins (fraction of disordered residues, melting temperature protein length) as described before^34^ and compared them across the same groups as above using a Mann-Whitney U test with Benjamini Hochberg correction (Supplementary Table 7).

### Upstream kinase investigation

The upstream kinases of phosphosites on various groups of proteins were evaluated using data from PhosphoSitePlus^98^ (version May 13, 2024), grouped into the kinase families reported in KinMap^99^ (accessed May 20, 2024). The counts of phosphosites for CCD proteins and enzymes were compared to those from the 13,147 proteins in the HPA Subcellular Atlas v23 to establish significance using the one-sided Fisher exact test, corrected by the Bonferroni method, and variable enzymes were similarly compared to all other enzymes (Supplementary Table 8). These results should be taken as qualitative assessments of PTM regulation, owing to the complexity of cellular regulation.

### Generation of clonal mNG-tagged cell lines

CRISPR/Cas9 mediated mNG tagging was performed to establish a polyclonal mNG tagged Hek293T cell lines as described previously.^16,100^ Afterwards, single cells were sorted into the wells of a 96 well plate and cultured in DMEM supplemented with 20% fetal bovine serum (FBS), 1 mM glutamine and 100 µg/ml penicillin/streptomycin (Gibco). to establish clonal populations. Cells were maintained in 96 well plates at 37 °C in a 5.0 % CO2 humidified environment and the medium was changed every three days until confluence was achieved. After 2-3 passages in complete DMEM with 10% FBS gDNA was extracted to confirm the genotype by Illumina amplicon sequencing and live cell microscopy was performed.

### Evaluating the genotype of mNG-tagged cell lines

DNA repair outcomes were characterized by Illumina amplicon sequencing. Clonally expanded Hek293T cells were grown in a 96-well plate at around 80% confluency, washed with PBS and resuspended in 50 μl QuickExtract (Lucigen) at 65 °C for 20 min and 98 °C for 5 min. Next, 2 μl gDNA, 20 μl 2× KAPA, 1.6 μl of 50 μM forward and reverse primer, 8 μl 5 M betaine and 8.4 μl H2O were run on a thermocycler for 3 min at 95 °C followed by 3 cycles of 20 s at 98 °C, 15 s at 63 °C, 20 s at 72 °C, 3 cycles of 20 s at 98 °C, 15 s at 65 °C, 20 s at 72 °C, 3 cycles of 20 s at 98 °C, 15 s at 67 °C, 20 s at 72 °C and 17 cycles of 20 s at 98 °C, 15 s at 69 °C, 20 s at 72 °C. Next, 1 μl of 2 nM PCR product, 4 μl of forward and reverse indexed barcoding primer, 20 μl 2× Kapa and 11 μl H2O were run with the following thermocycler settings: 3 min at 95 °C, 10 cycles of 20 s at 98 °C, 15 s at 68 °C, 12 s at 72 °C. Product concentrations in the 200–600 bp range were quantified using a fragment analyzer (Agilent) and pooled at 500 nM. The sequencing library was purified using dual solid phase reversible immobilization at a 0.6 and 1.1× bead/sample ratio. Sequencing was performed using an Illumina MiSeq system at the CZ Biohub Sequencing facility. Sequencing outcomes were characterized using CRISPResso^101^ (version 1.0.13). For clonal expansion and time lapse microscopy only clones with a homozygous homology directed repair (HDR) mediated knock in were considered.

### Spinning-disk confocal live cell microscopy

Approximately 20,000 endogenously tagged HEK293T cells were grown on a fibronectin (Roche)-coated 96 well glass bottom plate (Cellvis) for 24 hours. Cells were counterstained in 0.5 µg/ml Hoechst 33342 (Thermo) for 30 min at 37 °C and imaged in complete DMEM without phenol-red. Live-cell imaging was performed at 37 °C and 5% CO_2_ on a Dragonfly spinning-disk confocal microscope (Andor) equipped with a 1.45 N/A 63x oil objective and an iXon Ultra 888 EMCCD camera (Andor) using the Micromanager software v1.4.^102^ For time-lapse acquisition of HMGCS1-mNG cells approximately 10,000 endogenously tagged HEK293T cells were prepared for live cell imaging, as described earlier. Images were acquired in hourly intervals for 40 hours.

### Iterative indirect immunofluorescence imaging (4i)

U-2OS cells were grown in McCoys 5A medium supplemented with 10% FBS and 1% PenStrep in a 5% CO2, 37 °C environment. Cells were passaged at 80% confluency. 8000 cells were seeded in a well of a 96 well plate and cultured for 24 hours. Cells were washed with PBS, fixed in 4% paraformaldehyde for 15 minutes, washed again and permeabilized in 0.1% Triton X-100 for 3x 5 minutes. Next, 4i was performed as previously described^103^. In every cycle, cells were (1) blocked, (2) stained with primary antibodies, secondary antibodies and DAPI, (3) imaged, and then (4) the antibodies were eluted. (1) Cells were washed three times with PBS and blocked in 4i blocking buffer (100 mL PBS, 0.535 g NH4Cl, 1 g bovine serum albumin, 1.45 g maleimide) for 1 h at RT. (2) Cells were washed four times with PBS and incubated with the primary antibody solution for 2 h at RT (Extended Data Table 2, 3). Cells were washed four times with PBS and incubated with the secondary antibodies for 1.5 h at RT. Antibodies were diluted to target concentration in blocking solution (100 mL PBS, 0.535 g NH4Cl, 1 g bovine serum albumin) (Extended Data Table 2). Cells were washed four times with PBS, and nuclear DNA was stained with 18 μM DAPI in PBS. Then, cells were washed four times with PBS, sealed with 100 ul imaging buffer (25 mL mQ water, 2.9 g N-acetyl-cysteine, pH 7.4) and, (3) imaged as described below. (4) Cells were washed and antibodies were eluted for 3x 10 minutes in elution buffer (0.19 g L-glycine, 1.9 mL 8M urea, 1.9 mL 8M guanidinum chloride, 0.1 g TCEP, volume adjusted to 5 mL with mQ water, pH 2.5).

### 4i image acquisition

Fluorescence images of immunostained U-2 OS cells were acquired on Operetta Phenix High-Content Screening System (Revvity) with the 20x water-immersion objective (NA=1.0, WD=1.7 mm, FOV=650×650 μm). Images were acquired from 9 fields per well at fixed positions as a z-stack of 9 slices spaced at 0.8 μm. For each position, images were acquired using the following excitation wavelengths: 530-560 nm for the marker proteins (Alexa Fluor™ 555, emission filter 570-650 nm), 460-490 nm for microtubules (Alexa Fluor™ 488, emission filter 500-550 nm), 615-645 nm for the proteins of interest (Alexa Fluor™ 647, emission spectra 655-760 nm), and 355-385 nm for DAPI staining (emission filter 430-500 nm). For the target proteins, images were acquired using two different exposure times to accommodate for proteins with particularly high signal intensities. Pixel dimensions of all images are 2160 x 2160. Images were acquired after each staining and each elution round.

### 4i image processing and analysis

All image processing was done in Python, unless otherwise specified. For each field of view, maximum projection of the z-stack was calculated using ImageJ 1.54f. Images from the tubulin channel for each cycle were used to align the fields of view between the cycles. Tubulin and DAPI channels from the first cycle were used for image segmentation using a previously described DPN-UNet segmentation model^96^, and cell and nuclei masks were stored for future image processing. Each cell was assigned a unique identifier, including the well and the field of view. Illumination correction was first performed in CellProfiler, and corrected image intensities were normalized within the well by calculating the average and standard deviation of all pixels within 100 randomly selected cells across all fields of view and performing reverse z-scoring of cell pixels in each individual field of view with these values. Cell and nuclei masks were used to extract the following parameters for each cell:

- Whether the cell is completely within the field of view (= not touches the border of the image). Incomplete cells were excluded from the analysis;
- Cell area and nucleus area in pixels. Cells with nuclei smaller than 1500 pixels were excluded from the analysis based on the visual assessment of the images;
- Ratio between the area of the nucleus and the area of the cytoplasm;
- Signal intensities in each of the channels. Signal intensity was defined as the average of the values of the top 25% of the pixels. Signal intensity was calculated for the entire cell, for the nucleus and for the cytoplasm (cell mask excluding nucleus mask) separately. Ratio of signal intensity between the nucleus and the cytoplasm was also calculated.

Morphological and intensity features pertaining to DAPI and marker stainings (16 features per cell), z-scored across the entire dataset of 46,274 cells, were used to create the UMAP representation of the dataset. Statistical analysis of correlations between the features and data visualization was performed in Python 3.10 (pandas 2.1.4, scipy 1.11.4, umap 0.1.1, seaborn 0.13.2)

### Circos plots for visualization of patterns of multilocalization within cells

The locations for each protein were extracted from the HPA subcellular section and grouped into meta structures and compartments as described previously. Multilocalizing proteins were visualized by a connection between the respective locations in the Circos plot, with proteins localizing to three or more locations displaying multiple connections in the plot. Unilocalizing proteins do not display any connections in the plot. To analyze the degree of experimentally determined protein multilocalization (excluding information derived from the HPA, we extracted subcellular location data for the proteins included on HPA subcellular section v23 from UniProt^104^ (20230221). We filtered for locations with experimental evidence (ECO:0000269) and converted the UniProt locations to matching GO terms as described previously (Supplementary Table 10).^15^ Finally, the degree of multilocalization in UniProt data were visualized as described before.

### Interactomics analysis to identify potentially multifunctional enzymes

We identified a protein’s interaction partners from various resources^16,65,66^ as well as from the STRING database^67^ (experimental evidence and confidence > 0.7) (Supplementary Table 11). Subsequently, a cellular component (nucleus, cytosol, nucleus+cytosol) was assigned to each interactor according to its localization data in the HPA Subcellular Section. Finally, an interaction network among all interactors was constructed with the color of the nodes indicating the subcellular protein location and the thickness of the edges showing the confidence of an interaction between 1 (experimentally verified location from the interactomics datasets) and 0.7 (lowest confidence extracted from theString database) using the matplotlib and *networkx* packages in python 3.10. This approach allowed us to evaluate the spatially-resolved interaction network for each enzyme and manually assess its non-metabolic functions using a guilty-by-association approach.

## Acknowledgements

We acknowledge the entire staff of the HPA. Funding was provided by the Knut and Alice Wallenberg Foundation to the Human Protein Atlas and E.L. (KAW 2021.0346 & KAW 2021.0189), Göran Gustafsson Stiftelse, Vetenskapsrådet to E.L (VR 2017-05327) and Schmidt Futures to E.L. A.S. was supported by a Rubicon grant from the Dutch Research Council (NWO, 019.231EN.013).

## Author contributions

C.G. participated in the development of the idea, conducted the experiments, conducted the analysis, generated figures and co-wrote the manuscript. A.C. and A.S. co-wrote the manuscript. A.C. conducted upstream kinase analysis. A.S. performed 4i imaging and analysis. T.L. performed cosine distance analysis and quantitative analysis of single-cell heterogeneity. D.M. participated in manuscript writing and contributed to discussions. S.S. and R.S. performed tissue image annotation. P.R. performed live-cell time lapse experiments. M.L. provided resources for, aided and supervised the generation of clonal mNG-tagged cell lines. C.L. provided resources for tissue image annotation and contributed to discussions. M.U. provided antibody resources. U.A. and A.S. were responsible for data integration into the proteinatlas.org database. E.L. developed the idea, designed and supervised the project and co-wrote the manuscript. All authors have read and approved the manuscript.

## Data and materials availability

All cell types are authenticated and listed in the Materials and Methods. Images are publicly available on the HPA website (www.proteinatlas.org), and single-cell transcriptomic data are available in GEO SRA project GSE146773. Uncompressed images for the cell cycle-resolved imaging proteomic dataset were annotated using IDR metadata templates and deposited in the BioImage Archive (accession S-BIAD34, https://www.ebi.ac.uk/biostudies/BioImages/studies/S-BIAD34). All other imaging data will be made publicly available upon publication.

## Code availability

Code will be made publicly available upon publication.

